# Neural Activity in Afferent Projections to the Infralimbic Cortex is Associated with Individual Differences in the Recall of Fear Extinction

**DOI:** 10.1101/2022.03.12.484108

**Authors:** Amanda S. Russo, Ryan G. Parsons

**Affiliations:** Stony Brook University

## Abstract

Post-traumatic stress disorder (PTSD) is characterized by an impaired ability to extinguish fear responses to trauma-associated cues. Studies in humans and non-human animals point to differences in the engagement of certain frontal cortical regions as key mediators determining whether or not fear extinction is successful, however the neural circuit interactions that dictate the differential involvement of these regions are not well understood. To better understand how individual differences in extinction recall are reflected in differences in neural circuit activity, we labeled projections to the infralimbic cortex (IL) in rats using a retrograde tracer and compared neural activity within, and outside, of IL-projecting neurons. We analyzed these data in groups separated on the basis of how well rats retained extinction memory. We found that within IL-projecting cells, neurons in the posterior paraventricular thalamus showed heightened activity in rats that showed good extinction recall. Outside of the IL-projecting cells, increased Fos activity was observed in good extinction rats in select regions of the claustrum and ventral hippocampus. Our results indicate that differences in extinction recall are associated with a specific pattern of neural activity both within and outside of projections to the IL.

## Introduction

Fear conditioning occurs when a neutral stimulus becomes associated with an aversive, unconditioned stimulus (UCS) such that the originally neutral stimulus, now the conditioned stimulus (CS), elicits a conditioned fear response (CR) in the absence of the UCS. Extinction of conditioned fear is the reduction in the CR to the CS as a result of repeated presentations of the CS alone [1]. Prior work suggests that post-traumatic stress disorder (PTSD) involves the inability to recall the extinction of a fear response [2], however the neural mechanisms that distinguish successful extinction recall from extinction failure are not clear [3,4].

Rodent models are useful in this endeavor because there is marked individual variation in extinction recall among rodents [5-8]. Prior work investigating the neural mechanisms of fear extinction at the group level has suggested that activation of the infralimbic cortex (IL) is necessary for extinction recall [9-11, but see 12], and a handful of studies have found reduced activity in the IL among rodents displaying poor extinction recall compared to rodents that extinguish fear well [13-15,7]. However, the mechanisms underlying the differential engagement of the IL in rodents that extinguish fear readily versus those that show poor extinction are not well understood.

One possibility is that differences in fear extinction recall among individuals is the result of differential activation of specific afferents to the IL. Anatomical studies [16] have demonstrated that a variety of cortical and subcortical brain areas send dense projections to the IL, and in turn the IL sends efferent projections to a number of brain areas [17]. Group-level studies have found that IL projections to the amygdala are important for the acquisition of fear extinction [18-20], whereas inputs to the IL from basolateral amygdala (BLA) have been similarly implicated in extinction learning [21]. There are fewer studies pertaining to the involvement of IL-centered circuits in extinction recall, although recent work has implicated both the ventral and dorsal hippocampus [22,23,11] projections to the IL as being involved. Efferent projections of the IL to the thalamic nucleus reuniens also appear to be involved in the recall of fear extinction [24].

These prior studies are beginning to paint a picture of the neural circuit interactions involved in extinction recall, however there are very little data pertaining to whether or not activity in IL-centered neural circuits drive individual differences in extinction recall. Here, we sought to determine if differences in fear extinction recall across individuals are associated with changes in the activation of specific input to the IL. A viral GFP-conjugated retrograde tracer was infused into the IL of the rats prior to behavioral testing, and Fos activity in IL afferents after extinction recall, fear recall, and in rats given no behavioral testing was measured. Our results indicate that projections from the posterior portion of the paraventricular thalamus to the IL show heightened activity in rats which successfully recall extinction. Outside of IL projections, neural activity within specific regions of the claustrum and ventral hippocampus was increased in rats showing good extinction. Our results indicate that patterns of neural activity, both within and outside, of projections to the IL are associated with individual differences in fear extinction recall.

## Materials and Methods

### Subjects

Fifty-four adult, male, Sprague-Dawley rats (300-325 grams upon arrival) obtained from Charles River Laboratories (Raleigh, NC) served as subjects. The rats were housed in pairs, with food and water freely available, on a 12hour light/dark cycle (lights on at 7 am). Two cohorts of rats were used for these experiments (n = 28 and n = 26). After exclusions for death, surgical misses, failure to express GFP in target sites, poor tissue quality, and behavioral issues (explained throughout the methods), the extinction recall group contained 21 rats, the fear recall group contained 7 rats, and the home cage control group contained 7 rats (35 total rats included in final analyses). All procedures were approved by the Stony Brook University Institutional Animal Care and Use Committee and were in accordance with the National Institutes of Health guidelines for the care and use of laboratory animals.

### Surgery

Rats were handled for two days prior to surgery. Rats were anaesthetized with ketamine (87 mg/kg) and xylazine (10 mg/kg), and they were placed in a stereotaxic apparatus (Stoelting, Woodale, IL) and received a unilateral injection of AAVrg-CAG-GFP (Addgene, [25]) into the IL (left- and right-side injections were counterbalanced). In order to perform the injection, a 22-gauge cannula was lowered into place (AP: +3.00, ML: +/- 0.6, DV: −5.2). A 28-gauge internal cannula (connected to an infusion pump via PE 20 tubing) was inserted into the guide cannula and used to deliver 0.6 μL of the virus at a rate of 0.15 μL per minute and stayed in place for five minutes after the infusion was complete. After suturing rats were injected with Meloxicam (1mg/kg) and they were returned to their home cages once ambulatory. Rats stayed in their home cage for about 7 weeks to allow for recovery and retrograde transport of the virus. Three rats died while under anesthesia, leaving 51 rats (94%) which successfully recovered from surgery.

### Behavioral Apparatus

All procedures took place in 32 cm × 25 cm × 21 cm conditioning chambers (Clever Systems Inc., Reston, VA) which were located within sound attenuating 45.7-cm × 43.2-cm × 43.2-cm isolation boxes (Clever Sys. Inc.). During extinction training and extinction recall sessions, the context was altered to appear different than the original conditioning context. Context A (fear conditioning) contained 28-V, incandescent, house light bulbs (Chicago Miniature Lighting, UK), whereas Context B (extinction training, extinction recall testing, and fear recall testing) contained infrared LED lights (Univivi IR Illuminator, Shenzen, China; U48R). Furthermore, while Context A had shock grid floors with stainless steel and Plexiglas walls, Context B contained painted, metal inserts placed over the floor and walls. Context B was also altered in shape by placing a bent 33.5 × 21.3-cm metal insert inside the default conditioning chambers. In addition, in Context A, chambers were wiped down with 5% acetic acid, while in Context B chambers were wiped down with 5% ammonium hydroxide. Lastly, in Context B, rats were carried into the testing room in buckets as opposed to rolled in their home cages on carts. Overhead cameras recorded behavioral sessions, and the video signal in each chamber fed into software (FreezeScan 2.00, Clever Sys. Inc., Reston, VA) that scored freezing behavior based on pixel change. Parameters have been chosen such that computer-scored freezing behavior closely matches hand-scored behavior by a trained observer. Values indicating percent duration of time spent freezing were collapsed across 30 second bins.

### Behavioral Procedures

All behavioral procedures took place during the light portion of the light/dark cycle. Rats were handled for 5 days prior to the beginning of behavioral procedures and were carted into the behavioral room during the last three days of handling. On the first day of behavioral testing, an extinction recall group of rats was exposed to fear conditioning where they were placed into Context A, received a 6-minute habituation period where no stimuli were presented, and then received two combinations of a 4 kHz, 76 dB, 30 second tone and a co-terminating, 1.0 mA, 1 second foot-shock (2 min ITI). In all behavioral sessions, rats were returned to their home cages 2 minutes following the final stimulus presentation. The following day, rats in the extinction recall group were placed into the Context B chamber and were exposed to 20 presentations of the tone (2 min ITI) as extinction training following a 6-minute habituation period. The next day, rats in the extinction recall group were exposed to 4 presentations of the tone in Context B as an extinction recall test following a 6-minute habituation period. 60 minutes following the end of the behavioral session, rats in the extinction recall group were perfused. One group of fear recall control rats was exposed to the same procedure during the first day of fear conditioning in Context A. 48 hours later, these rats were placed into the Context B chamber and were exposed to 4 presentations of the tone (2 min ITI) as a fear recall test following a 6-minute habituation period. 60 minutes following the end of the behavioral session, these rats were perfused. One group of home cage control rats remained in their home cages throughout the duration of the experiment and were perfused on the same day as the experimental rats. Each of the two cohorts of rats was split into two runs, and the number of animals in each group was balanced across runs. One rat from the fear recall group was excluded from analyses because it failed to express evidence of fear conditioning (froze less than 15% of the time during the fear recall test). See schematic representation of the behavioral timeline in Figure 2A.

### Immunofluorescence

Rats were overdosed with Fatal Plus Solution (100 mg/kg) and perfused with ice-cold 10% PBS followed by 10% buffered formalin. The brains were extracted and stored in a solution of 30% sucrose-formalin at 4°C for approximately 1 week. Brains were then frozen and sectioned on a cryostat, and slices were taken at 40 μm thickness. Sections were stored in a serial order in 10% PBS at 4°C. Immunofluorescence was then performed on free-floating sections containing the brain areas of interest. Sections were washed 3 times in 10% PBS for 5 minutes each. Next, sections were incubated in 5% normal goat serum blocking solution for 2 hours at room temperature followed by 3 additional 5-minute-long washes in 10% PBS. The sections were then incubated in primary antibody (c-Fos, #2250, 1:500) (Cell Signaling, Danvers, MA) diluted in a solution of 1% BSA in 10% PBS at 4°C overnight. The next day, the sections were washed in 10% PBS at 4°C for 30 minutes, followed by 3, 5-minute-long washes with 10% PBS, and incubation in secondary antibody (Alexa Fluor 594 goat anti-rabbit, red conjugate, 1:500) (Invitrogen, Carlsbad, CA) at room temperature for 2 hours. After 3 more 5-minute-long washes in 10% PBS, the sections were mounted onto slides and cover-slipped with Fluoromount-G (Invitrogen). See representative images of the immunostaining in Figure 3G.

### Image Acquisition

Fluorescence microscopy using an Infinity3 digital camera (Lumenera, Ottawa, ON, Canada) with a light engine (Lumencor, Beaverton, OR) attached to a Zeiss microscope was used to acquire images from each brain area of interest, including slices containing the IL which did not undergo immunofluorescence, to confirm the correct placement of the injection site. Images used for cell counting were taken at 20x magnification. For each slice of tissue, one image was taken using a filter allowing visualization of GFP, one image was taken using a filter allowing visualization of the red conjugate of Alexa Fluor in the secondary antibody, and an imaging acquisition software (Infinity Analyze, Version 3) was used to overlay the images. All images across all brain areas were acquired with the same exposure time and gain setting. Six rats were excluded from analyses because the primary spread of the virus was outside of the IL (88% hit rate). Eight additional rats were excluded because although the virus hit the IL, they failed to show sufficient expression of GFP in all of the target brain areas of interest. Furthermore, one rat was excluded because of poor tissue quality.

### Cell Counting

Brightness and contrast were adjusted to reduce background noise in Image J (NIH) using an identical procedure for each image. Cell counts for total number of retrogradely-labeled cells, total number of Fos-labeled, and total number of double-labeled cells were performed manually using Image J’s “Cell Counter” plugin by an experimenter who was blinded to the identity of the animals. Cell counts were normalized to cells/mm^2^. For the analysis of Fos expression in IL-projecting cells, the number of double-labeled cells was normalized to the total number of retrogradely-labeled cells. For analysis of the mBLA, mvHPC, and pvHPC, cell counts from multiple 20x images were added together and normalized to cells/mm^2^. For analysis of the rest of the brain areas, one 20x image or a sub-portion of one 20x image was analyzed and normalized to cells/mm^2^. Analysis of the vHPC included the CA1, CA2, and subiculum areas of the vHPC. Figure 1 depicts the brain regions analyzed with images depicting boundaries across the anterior-posterior plane.

**Figure 1.**
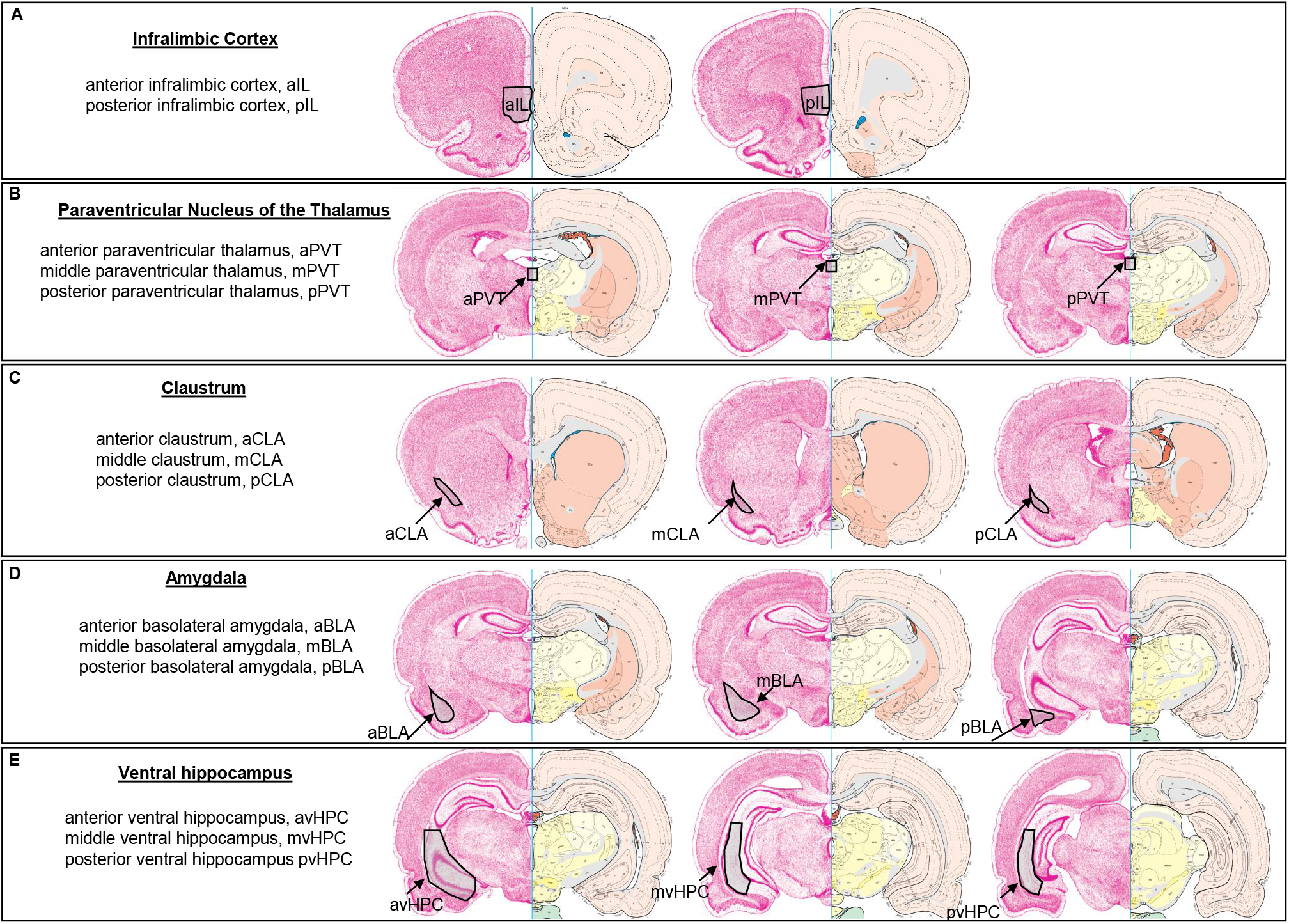
Abbreviations and Locations of Brain Areas of Interest. Explanation of abbreviations and locations for brain areas referenced throughout the manuscript.

### Experimental Design and Statistical Analysis

Extinction recall scores were calculated by expressing the percent time freezing during extinction recall as a percentage of freezing during the first 4 trials of extinction training. Low scores indicated good extinction recall, whereas high scores indicated poor extinction recall. The rats’ extinction recall scores were rank-ordered, and the rats which fell within the top third of extinction recall scores were classified as “poor extinction rats”, whereas the rats which fell within the bottom two thirds of extinction recall scores were classified as “good extinction recall rats.”

Non-parametric tests were used because data often violated assumptions of normal distribution and/or homogeneity of variance. Spearman’s rank order correlations were used to determine if there was a significant association between extinction recall scores and both Fos labeling and double-labeling in the brain areas of interest. Mann-Whitney U tests were used to determine whether there were differences between two independent groups. Kruskal-Wallis tests were used to determine whether 2 or more groups were different from one another, and Dunn’s multiple comparisons tests were used when the Kruskal-Wallis statistic was significant. A repeated-measures ANOVA with group as a between-subjects factor and trial as a within-subjects factor was used to assess freezing during extinction training. Results were considered significant when *p* < 0.05 for all statistical tests.

### Behavioral Results

Figure 2 depicts the experimental timeline (Figure 2A), and a frequency distribution for all rats that underwent extinction (Figure 2B). Rats in the good and poor extinction groups differed significantly in these calculated extinction recall scores (*U* = 0, *p* < 0.001) (Figure 2C). There was no significant difference among the good extinction, poor extinction, and fear recall groups in time spent freezing during the baseline period of the fear conditioning session (X^2^ = 2.746, *p* = 0.253) (Figure 2D). Furthermore, there was neither a significant difference among the good extinction, poor extinction, and fear recall groups in time spent freezing during the first tone presentation of the fear conditioning session (X^2^ = 1.107, *p* = 0.575), nor was there a significant difference among the good extinction, poor extinction, and fear recall groups in time spent freezing during the second tone presentation of the fear conditioning session (X^2^ = 2.214, *p* = 0.331) (Figure 2D). There was also no significant difference between the good and poor extinction groups in time spent freezing during the baseline period of the extinction training session (*U* = 45.00, *p* = 0.799) (Figure 2D). Next, there was a significant main effect of trial block (5 tones per block) on time spent freezing during the extinction training session (F (2.884, 54.80) = 8.331, *p* < 0.001), indicating that extinction learning occurred (Figure 2D). However, there was no main effect of extinction group (F (1, 19) = 3.091, *p* = 0.095) on time spent freezing throughout the extinction training session, nor was there an interaction between trial block and extinction group (F (4, 76) = 1.890, *p* = 0.121) (Figure 2D). During the testing session, there was a significant difference among the good extinction, poor extinction, and fear recall groups in time spent freezing during the baseline period (X^2^ = 8.569, *p* = 0.014) such that the fear recall group froze significantly more than the good extinction group (*Mean Rank Diff*. = 10.57, *p* = 0.017), but not the poor extinction group (*Mean Rank Diff*. = −3.714, *p* > 0.999) (Figure 2D).

**Figure 2.**
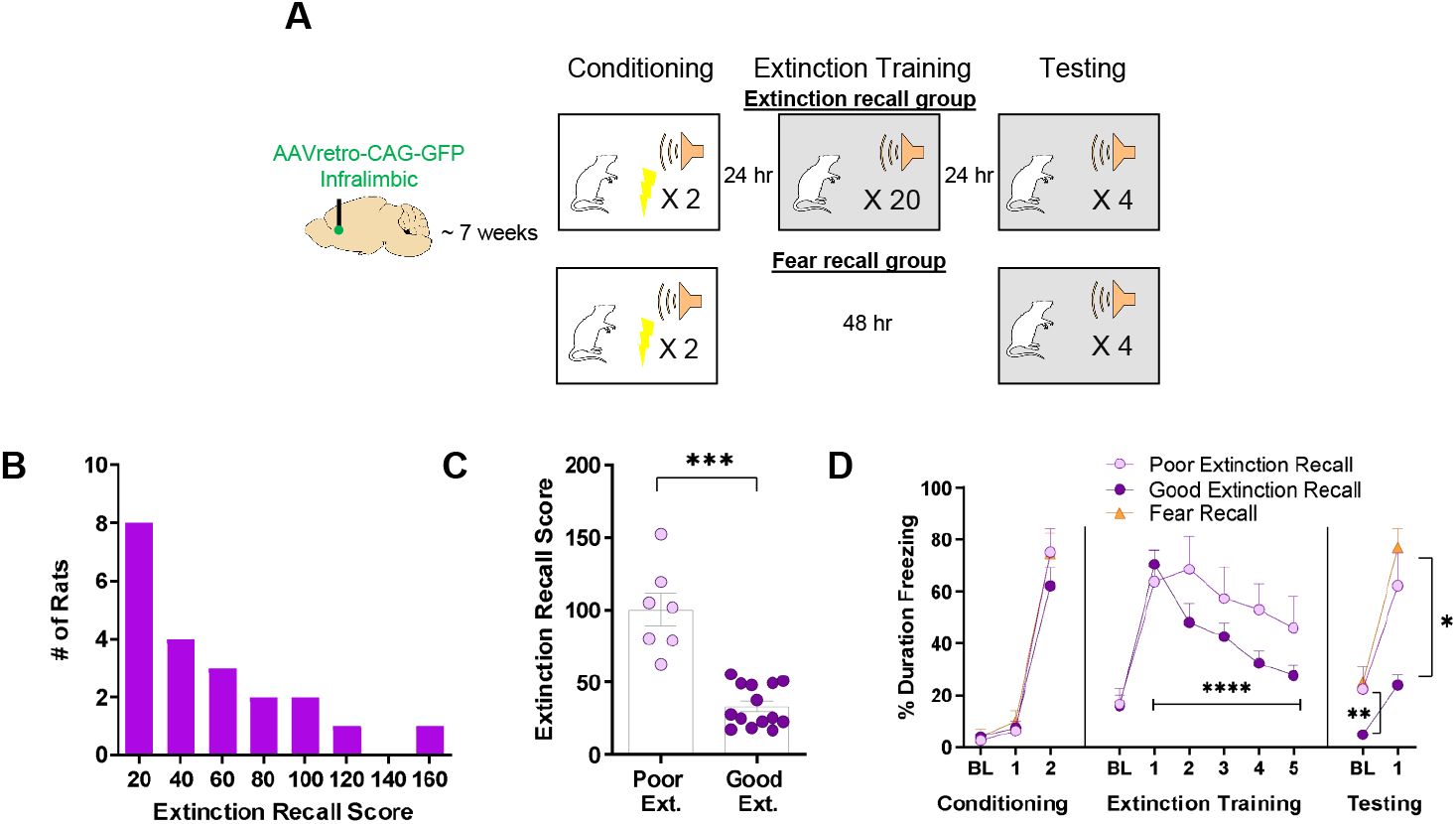
Individual Variation in Extinction Recall. (A) Schematic representation of surgical and behavioral procedures. (B) Frequency distribution demonstrating individual variation in extinction recall scores. (C) Evidence that groups created on the basis of calculated extinction recall scores represent two distinct phenotypes. (D) Percent time spent freezing for poor extinction, good extinction, and fear recall rats throughout behavioral procedures. Error bars represent standard error of the mean. * = *p* < 0.05, ** = *p* < 0.01, *** = *p* < 0.001, **** = *p* < 0.0001.

The good extinction, poor extinction, and fear recall groups also differed significantly from one another in time spent freezing during the tone presentations of the testing session (X^2^ = 14.93, *p* = 0.001) such that the good extinction group froze significantly less than both the poor extinction group (*Mean Rank Diff*. = 9.286, *p* = 0.044) and the fear recall group (*Mean Rank Diff*. = 13.86, *p* = 0.001) (Figure 2D).

### Anatomical Results

A retrograde tracer was infused into the IL (Figure 3A), and quantitative analysis was performed on the number of GFP+ cells along the anteroposterior axis of regions of interest (Figure 3B-F). There was a significant difference in the number of GFP+ cells among the anterior, middle, and posterior PVT (X^2^ = 8.200, *p* = 0.017) such that the mPVT displayed significantly more GFP+ cells than both the aPVT (*Mean Rank Diff*. = 18.37, *p* = 0.035) and the pPVT (*Mean Rank Diff*. = 17.71, *p* = 0.045) (Figure 3C). Although multiple animals did not display any GFP+ cells in the pCLA, yielding activity mapping in this area unfeasible, there was no significant difference among the anterior, middle, and posterior CLA (X^2^ = 5.596, *p* = 0.061) in the number of GFP+ cells (Figure 3D). Next, because very few rats displayed any GFP+ cells in the aBLA or the avHPC, only the middle and posterior portions of these regions were analyzed. There was a significant difference between the middle and posterior BLA (*U* = 393, *p* = 0.009) in the number of GFP+ cells such that the pBLA displayed more IL-projections than the mBLA (Figure 3E). Similarly, there was a significant difference between the middle and posterior vHPC such that the pvHPC displayed more IL-projections than the mvHPC (*U* = 403.5, *p* = 0.014) (Figure 3F).

**Figure 3.**
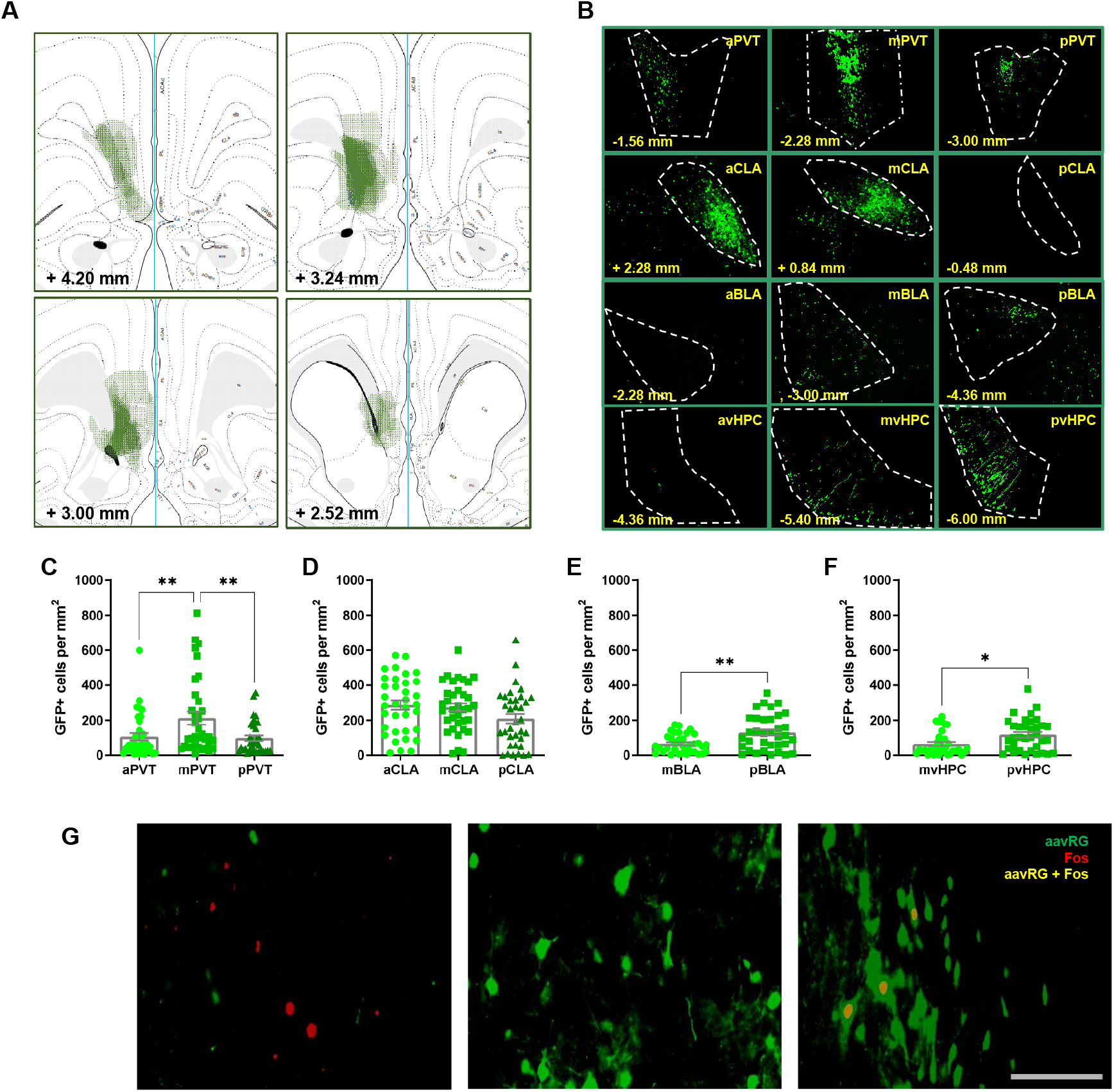
Quantification of IL-afferents throughout the Brain Areas of Interest. (A) Schematic representation of the aavRG-CAG-GFP spread throughout the IL of included rats. (B) Representative images of retrograde labeling in different anteroposterior locations of brain areas of interest. Quantification of retrograde label along the anteroposterior axis of the (C) paraventricular thalamus, (D) claustrum, (E) basolateral amygdala, and (F) ventral hippocampus. (G) Representative images demonstrating aavRG retrograde labeling, Fos labeling, and double-labeling of aavRG and Fos. Error bars represent standard error of the mean. * = *p* < 0.05, ** = *p* < 0.01. Scale bar = 100 µm.

### Activity Mapping Results

#### a. The Paraventricular Nucleus of the Thalamus

Both global and IL-projection specific Fos activity were analyzed in the aPVT, the mPVT, and the pPVT in all rats. There was no significant difference among the good extinction, poor extinction, fear recall, and home cage groups in Fos expression in the aPVT (X^2^ = 3.888, *p* = 0.274) (Figure 4A), nor was there a significant correlation between Fos expression in the aPVT and extinction recall (rs = 0.092, *p* = 0.691) (Figure 4B) or between Fos expression within IL-afferents of the aPVT and extinction recall (rs = 0.143, *p* = 0.537) (Figure 4D). However, there was a significant difference among good extinction, poor extinction, fear recall, and home cage groups in Fos expression within IL-afferents of the aPVT (X^2^ = 15.05, *p* = 0.002) such that the fear recall group displayed more activation of IL-afferents relative to the good extinction (*Mean Rank Diff*. = 11.54, *p* = 0.003), poor extinction (*Mean Rank Diff*. = 10.57, *p* = 0.034), and home cage (*Mean Rank Diff*. = 12.79, *p* = 0.005) groups (Figure 4C). Next, there was no significant difference among the good extinction, poor extinction, fear recall, and home cage groups in Fos expression in the mPVT (X^2^ = 2.272, *p* = 0.518) (Figure 4E), nor was there a significant correlation between Fos expression in the mPVT and extinction recall (rs = 0.168 *p* = 0.468) (Figure 4F). Although there was a significant difference among the good extinction, poor extinction, fear recall, and home cage groups in Fos expression within IL-afferents of the mPVT (X^2^ = 9.252, *p* = 0.026), post hoc comparisons did not reveal significant differences between any two groups (Figure 4G). Furthermore, there was no significant correlation between Fos expression within IL-afferents of the mPVT and extinction recall (rs = 0.174, *p* = 0.450) (Figure 4H). Next, there was a significant difference among the good extinction, poor extinction, fear recall, and home cage groups in Fos expression in the pPVT (X^2^ = 13.89, *p* = 0.003), such that the good extinction group (*Mean Rank Diff*. = 14.96, *p* = 0.010), but not the poor extinction (*Mean Rank Diff*. = 12.86, *p* = 0.113) or fear recall group (*Mean Rank Diff*. = 2.571, *p* > 0.999), displayed more Fos expression than the home cage group (Figure 4I). There was, however, no significant correlation between Fos expression in the pPVT and extinction recall (rs = 0.051, *p* = 0.825) (Figure 4J). Finally, there was both a significant difference among the good extinction, poor extinction, fear recall, and home cage groups in Fos expression within IL-afferents of the pPVT (X^2^ = 12.34 *p* = 0.006) such that the good extinction group displayed more Fos expression in IL-afferents than both the poor extinction (*Mean Rank Diff*. = 12.54, *p* = 0.014) and home cage (*Mean Rank Diff*. = 12.89, *p* = 0.049) groups (Figure 4K) and a significant correlation between activation of IL-afferents within the pPVT and extinction recall such that better extinction recall was related to more activation of these IL-afferents (rs = −0.438, *p* = 0.047) (Figure 4L).

**Figure 4.**
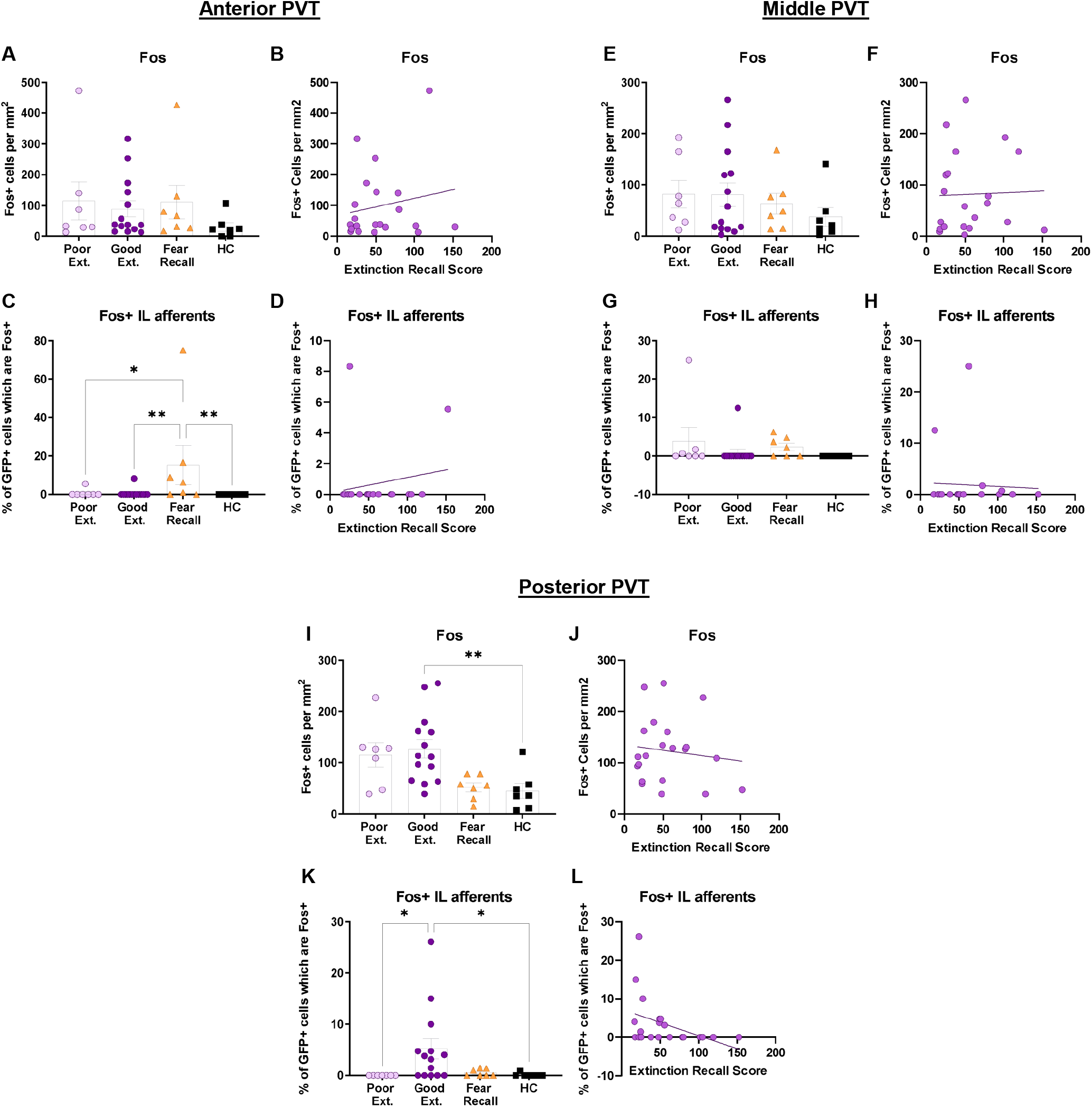
Fos Activity is Elevated in the IL-afferents of the Posterior Paraventricular Thalamus (PVT) in Rats Displaying Good Extinction. (A) No significant differences among groups in Fos expression in the aPVT. (B) No significant correlation between Fos expression and extinction recall in the aPVT. (C) Fear recall group showed elevated Fos expression within IL-afferents relative to all other groups. (D) No significant correlation between Fos expression within IL-afferents and extinction recall in the aPVT. (E) No significant differences among groups in Fos expression in the mPVT. (F) No significant correlation between Fos expression and extinction recall in the mPVT. (G) No significant differences among groups in Fos expression within IL-afferents in the mPVT. (H) No significant correlation between Fos expression within IL-afferents and extinction recall in the mPVT. (I) Good extinction group, but no other groups, shows elevated Fos activity in the pPVT relative to the home cage group. (J) No significant correlation between Fos expression and extinction recall in the pPVT. (K) Good extinction group shows elevated Fos expression in IL-afferents relative to the poor extinction group and the home cage group. (L) Significant correlation between Fos expression within IL-afferents and extinction recall such that good extinction recall is related to more Fos expression within IL-afferents.. Error bars represent standard error of the mean. * = *p* < 0.05, ** = *p* < 0.01.

#### b. The Claustrum

Next, global and IL-projection specific Fos activity were analyzed in the aCLA and mCLA in rats in all groups. There was a significant difference among the good extinction, poor extinction, fear recall, and home cage groups in Fos expression in the aCLA (X^2^ = 8.455, *p* = 0.036) such that the fear recall group (*Mean Rank Diff*. = 14.50, *p* = 0.049), but neither the poor (*Mean Rank Diff*. = 10.21, *p* = 0.373) nor good extinction (*Mean Rank Diff*. = 4.607, *p* > 0.999) groups, displayed more Fos expression than the home cage group (Figure 5A). There was no significant correlation between global Fos expression in (rs = 0.036, *p* = 0.876) (Figure 5B) or Fos expression within (rs = −0.282, *p* = 0.215) the IL-afferents of the aCLA and extinction recall (Figure 5D), nor was there a significant difference among the good extinction, poor extinction, fear recall, and home cage groups in Fos expression within the IL-afferents of the aCLA (X^2^ = 6.722, *p* = 0.081) (Figure 5C). Next, there was a significant difference among the good extinction, poor extinction, fear recall, and home cage groups in Fos expression in the mCLA (X^2^ = 10.12, *p* = 0.018) such that the good extinction group (*Mean Rank Diff*. = 12.93, *p* = 0.038), but neither the poor extinction (*Mean Rank Diff*. = 5.143, *p* > 0.999) nor fear recall groups (*Mean Rank Diff*. = 14.00, *p* = 0.063) displayed significantly more Fos expression in the mCLA relative to the home cage group (Figure 5E). However, there was no significant correlation between global Fos expression in the mCLA (rs = 0.321, *p* = 0.156) (Figure 5F) or Fos expression within the IL-afferents of the mCLA (rs = −0.121, *p* = 0.602) and extinction recall (Figure 5H), nor was there a significant difference among the good extinction, poor extinction, fear recall, and home cage groups in Fos expression within the IL-afferents of the mCLA (X^2^ = 4.923, *p* = 0.178) (Figure 5G).

**Figure 5.**
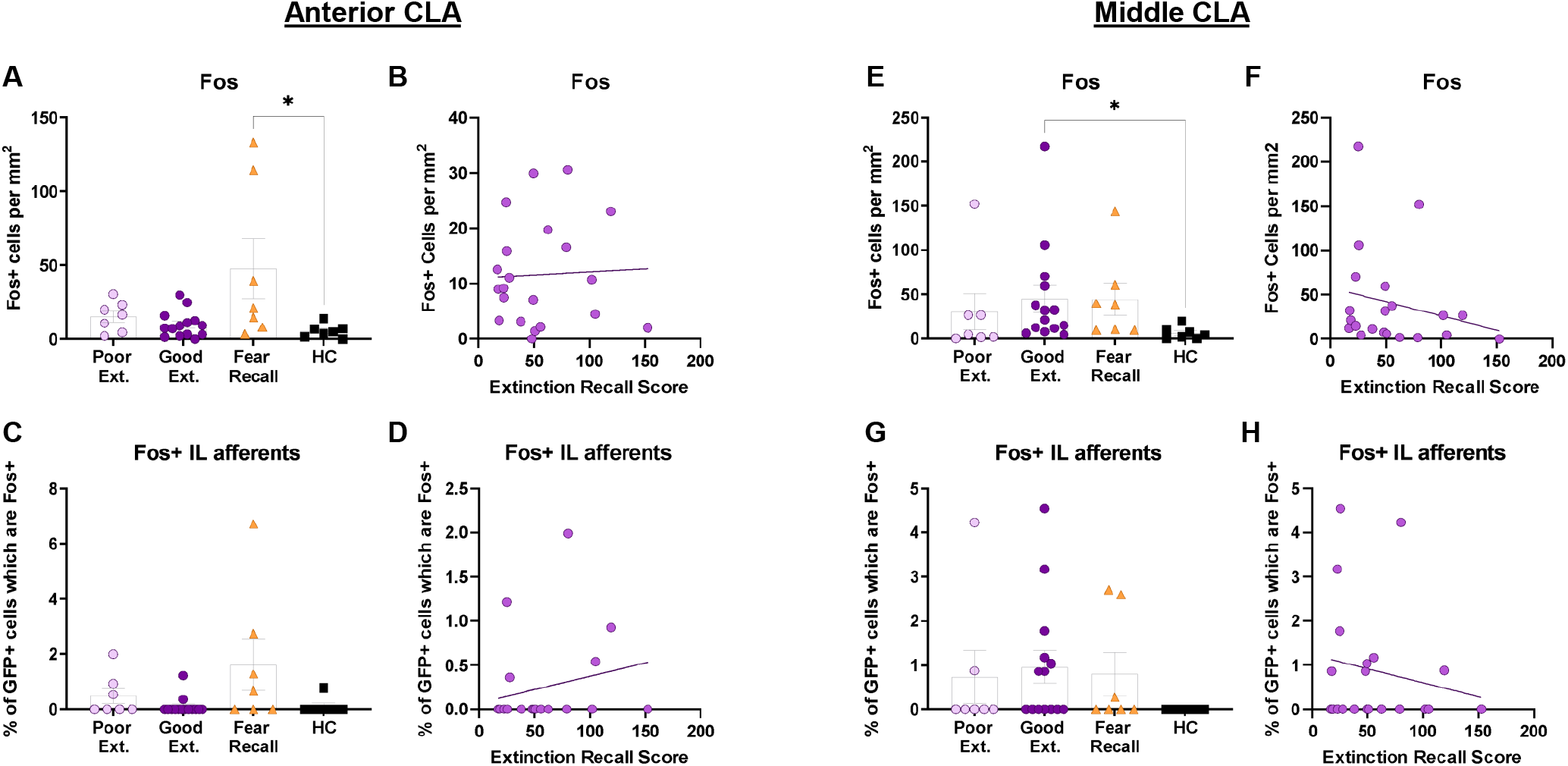
Fos Activity is Elevated in the Middle Claustrum among Rats Displaying Good Extinction Recall. (A) Fear recall group, but not other group, shows elevated Fos activity relative to the home cage group in the aCLA. (B) No significant correlation between Fos expression in the aCLA and extinction recall. (C) No significant difference among groups in Fos expression within the IL-afferents of the aCLA. (D) No significant correlation between Fos expression within the IL-afferents and extinction recall in the aCLA. (E) Good extinction group, but not other group, shows elevated Fos activity relative to the home cage group in the mCLA. (F) No significant correlation between Fos expression and extinction recall in the mCLA. (G) No significant differences among groups in Fos expression within the IL-afferents of the mCLA. (H) No significant correlation between Fos expression within the IL-afferents and extinction recall in the mCLA. Error bars represent standard error of the mean. * = *p* < 0.05.

#### c. The Basolateral Amygdala

Next, global and IL-projection specific Fos activity were analyzed in the mBLA and the pBLA in rats in all groups. There was no significant difference among the good extinction, poor extinction, fear recall, and home cage groups in Fos expression in the mBLA (X^2^ = 0.944, *p* = 0.815) (Figure 6A). There was also no significant difference among the good extinction, poor extinction, fear recall, and home cage groups in Fos expression in the IL-afferents of the mBLA (X^2^ = 0.518, *p* = 0.915) (Figure 6C). Furthermore, there was neither a significant correlation between global Fos expression in the mBLA (rs = 0.126, *p* = 0.588) (Figure 6B) nor Fos expression within the IL-afferents of the mBLA (rs = 0.200, *p* = 0.385) (Figure 6D) and extinction recall. There was also no significant difference among the good extinction, poor extinction, fear recall, and home cage groups in Fos expression in the pBLA (X^2^ = 4.246, *p* = 0.236) (Figure 6E), and no significant difference among the good extinction, poor extinction, fear recall, and home cage groups in Fos expression within the IL-afferents of the pBLA (X^2^ = 1.954, *p* = 0.582) (Figure 6G). Finally, there was neither a significant correlation between global Fos expression in the pBLA (rs = 0.070, *p* = 0.762) (Figure 6F) nor Fos expression within the IL-afferents of the pBLA (rs = 0.122, *p* = 0.597) and extinction recall (Figure 6H).

**Figure 6.**
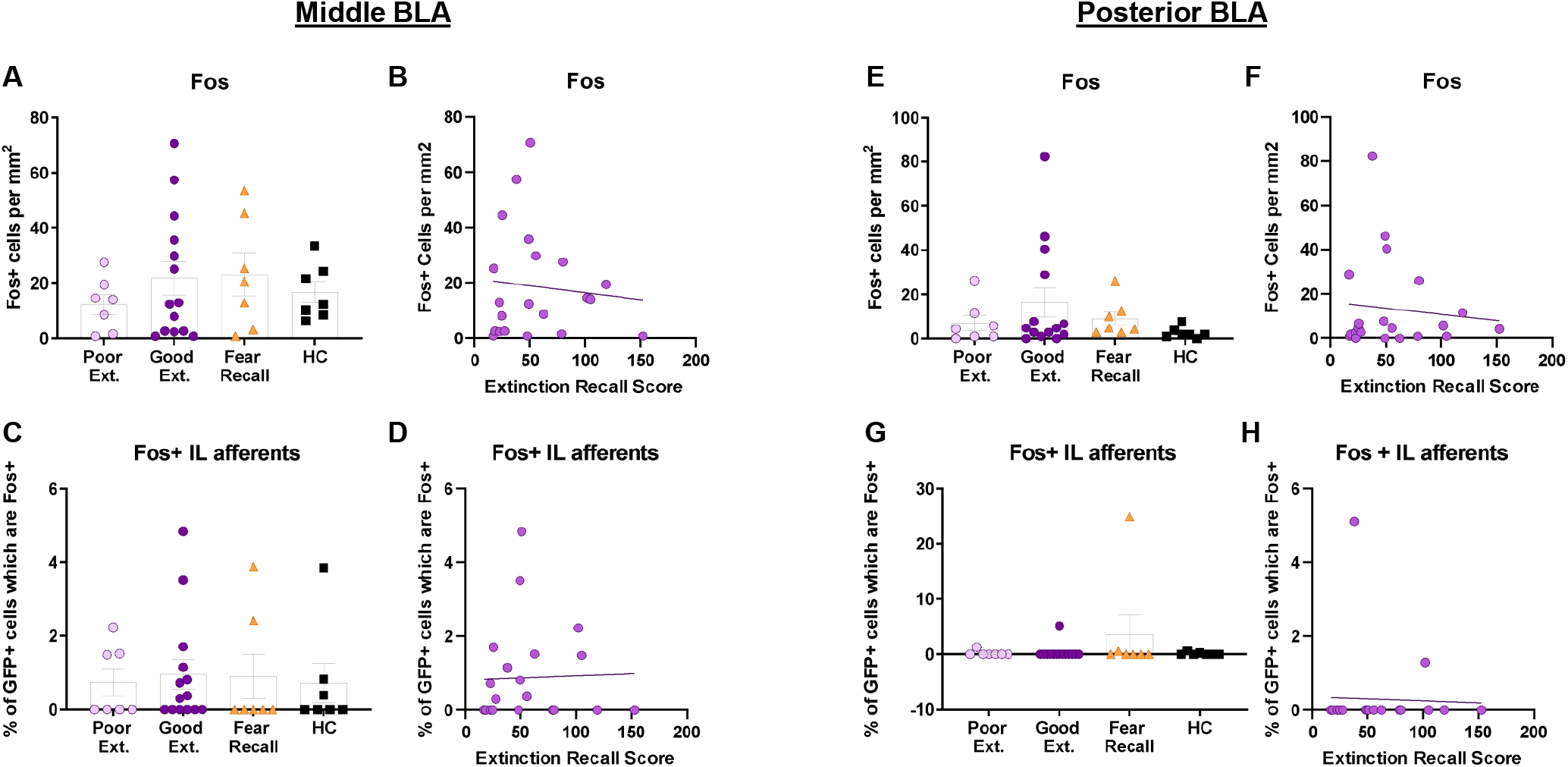
Individual Variation in Extinction Recall Does Not Map onto Differences in Fos Expression in the Basolateral Amygdala. (A) No significant differences among groups in Fos expression in the mBLA. (B) No significant correlation between Fos expression and extinction recall in the mBLA. (C) No significant differences among groups in Fos expression within the IL-afferents of the mBLA. (D) No significant correlation between Fos expression within the IL-afferents and extinction recall in the mBLA. (E) No significant differences among groups in Fos expression in the pBLA. (F) No significant correlation between Fos expression and extinction recall in the pBLA. (G) No significant differences among groups in Fos-expression within IL-afferents of the pBLA. (H) No significant correlation between Fos expression within IL-afferents and extinction recall in the pBLA. Error bars represent standard error of the mean.

#### d. The Ventral Hippocampus

Finally, global and IL-projection specific Fos activity were analyzed in the mvHPC and the pvHPC in all rats. There was a significant difference among the good extinction, poor extinction, fear recall, and home cage groups in Fos expression in the mvHPC (X^2^ = 8.056, *p* = 0.045) such that the good extinction (*Mean Rank Diff*. = 13.29, *p* = 0.031), but neither poor extinction (*Mean Rank Diff*. = 6.857, *p* > 0.999) nor fear recall (*Mean Rank Diff*. = 8.000, *p* = 0.864) groups showed more Fos expression than the home cage group (Figure 7A). There was, however, no significant difference among the good extinction, poor extinction, fear recall, and home cage (*Mdn* = 0.000, *Range* = 0.433) groups in Fos expression in the IL-afferents of the mvHPC (X^2^ = 4.893, *p* = 0.180) (Figure 7C). Furthermore, there was neither a significant correlation between global Fos expression in the mvHPC (rs = −0.233, *p* = 0.309) (Figure 7B) nor Fos expression within the IL-afferents of the mvHPC (rs = 0.056, *p* = 0.810) (Figure 7D) and extinction recall. Next, there was no significant difference among the good extinction, poor extinction, fear recall, and home cage groups in Fos expression in the pvHPC (X^2^ = 3.623, *p* = 0.353) (Figure 7E), and no significant difference among the good extinction, poor extinction, fear recall, and home cage groups in Fos expression within the IL-afferents of the pvHPC (X^2^ = 3.871, *p* = 0.276) (Figure 7G). Finally, there was neither a significant correlation between global Fos expression in the pvHPC (rs = −0.127, *p* = 0.584) (Figure 7F) nor Fos expression within the IL-afferents of the pBLA (rs = 0.176, *p* = 0.447) and extinction recall (Figure 7H).

**Figure 7.**
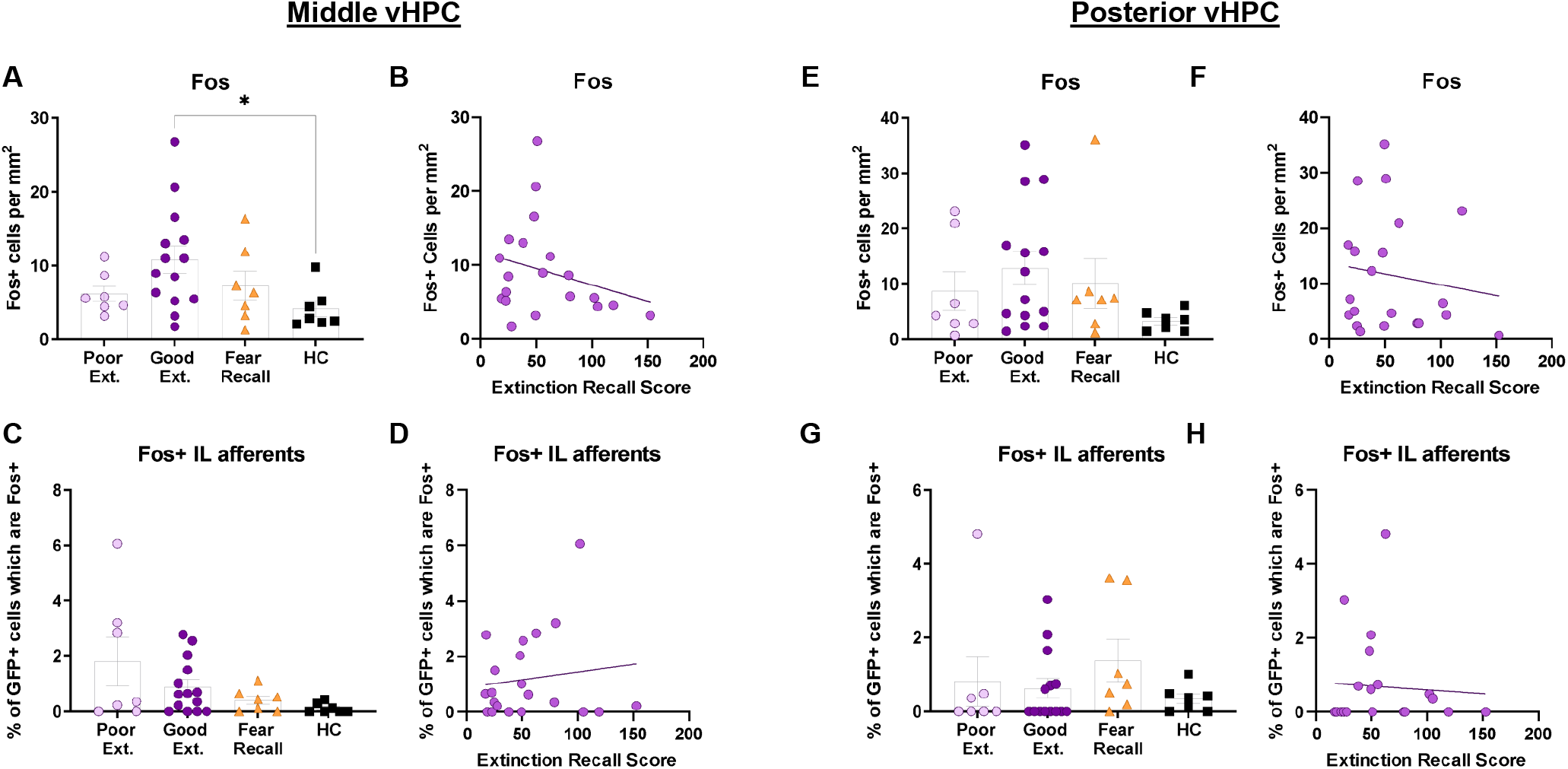
Elevated Fos Expression in the Middle Ventral Hippocampus of Rats Displaying Good Extinction Recall. (A) Good extinction group, but not other group, shows elevated Fos expression relative to the home cage group in the mvHPC. (B) No significant correlation between Fos expression and extinction recall in the mvPHC. (C) No significant differences among groups in Fos expression within IL-afferents of the mvHPC. (D) No significant correlation between Fos expression within IL-afferents and extinction recall in the mvHPC. (E) No significant differences among groups in Fos expression in the pvHPC. (F) No significant correlation between Fos expression and extinction recall in the pvHPC. (G) No significant differences among groups in Fos expression within the IL-afferents of the pVHPC. (H) No significant correlation between Fos expression within IL-afferents and extinction recall in the pvHPC. Error bars represent standard error of the mean. * = *p* < 0.05.

## Discussion

Here, we tested whether individual differences in extinction recall would be reflected in different patterns of activity in afferents to the infralimbic cortex. To this end, we assessed Fos activity in projections to the IL from the paraventricular thalamus, claustrum, basolateral amygdala, and ventral hippocampus following extinction recall. Within cells that project to the IL, we found that activity in the posterior region of the PVT was higher in rats which showed good extinction recall compared to rats with poor extinction. No differences were observed in IL afferents from the claustrum, ventral hippocampus, or basolateral amygdala. Outside of the IL-projecting cells, increased neural activity was observed in good extinction rats in select regions of the claustrum and ventral hippocampus. Our results suggest that successful extinction recall is orchestrated by specific PVT projections to the IL and non-IL targeting cells in the claustrum and ventral hippocampus.

Our finding that PVT projections to the IL are active in rats that showed good extinction recall is consistent with a recent study showing that the PVT is necessary for extinction recall [26]. This study did not employ manipulations that were sub-region specific, however they did show that both PVT projections to the lateral division of the central amygdala, and IL projections to the PVT, are necessary for extinction recall. Our results suggest that in addition to the IL-PVT-CeL circuit, posterior PVT inputs to the IL might also be required for extinction recall. Thus, it appears as though both efferent and afferent connections of the IL are involved in extinction recall. An important next step will be to determine what drives the pPVT to signal extinction recall at the neural circuit level. In addition to reciprocal connections to the IL, prior tract tracing studies [27,28] have showed that the pPVT receives input from the ventral periaqueductal gray (vPAG), which has been implicated in extinction learning [29-32]. While the role of the vPAG in extinction recall has not been established, projections to the pPVT from the vPAG are an attractive candidate given their density and prior evidence of both regions’ involvement in the acquisition of fear extinction.

Another important aspect of our results from the PVT is that they are specialized along its anteroposterior axis. Strikingly, neural activity within PVT projections to IL were associated with opposing behavioral states such that activity in anterior PVT projections to the IL were associated with fear recall, whereas pPVT projections were active following successful recall of extinction (i.e. low fear). Such heterogeneity of function within the PVT is not surprising given prior work [discussed in 33]. A prime example of the distribution of function within the PVT came recently [34] in a study which characterized the properties of specific cell types in the PVT. This study showed that DRD2-expressing dopamine cells, preferentially expressed in the pPVT, innervated the prelimbic cortex and were responsive to aversive stimuli. A second population of cells were predominantly expressed in the aPVT and signaled a transition into a state of low physiological arousal, and which innervated the infralimbic cortex. Our results do not easily fit this framework, as aPVT cells projecting to the IL were active during fear recall and pPVT projections were active while animals were expressing low levels of fear. There are at least two possible explanations as to the apparent discrepancy. First, the identified cell types are not exclusively located in one anteroposterior location in the PVT. Thus, it is possible that the IL-projecting pPVT cells which are active in rats displaying good extinction recall belong to the class of cells which is more likely to be found in aPVT and signal a transition to a low arousal state. The same might be true for the IL-projecting cells in the aPVT which are activated following fear recall. Second, prior tracing studies have established the presence of pPVT projections to the IL [35], and while few appear to originate from DRD2-containing cells, it is possible that other cell types project to the IL and which are activated by successful extinction recall.

Although the goal of the present study was to identify differences among rats displaying different extinction phenotypes, these experiments also revealed novel findings pertaining to the mechanisms underlying fear recall. Of interest, we showed elevated Fos activity in the anterior portion of the CLA in rats expressing a fear memory. The claustrum is positioned as a hub for cortical connectivity and has been implicated in processes ranging from sensory integration, to attention and sleep [36-39]. There is limited data as to how the claustrum might participate in fear conditioning or the expression of fear, however, an earlier study showed that the expression of contextual fear engages Fos activity in the claustrum [40]. More recently, it was reported that inhibiting claustral projections to the entorhinal cortex during contextual fear conditioning attenuated long-term memory formation [41], although their requirement for the expression of fear was not tested. In the same study, increased Fos activation was observed when animals were exposed to a novel context compared to mice exposed to a familiar context. Given this, it is possible that the activation of CLA we report here is driven by the exposure to the novel chamber during testing and not fear recall per se. To more precisely understand the function of the claustrum in fear and contextual processing, future studies should employ targeted manipulations of claustrum.

Despite prior work implicating the PVT in fear memory expression [42-44], we did not observe any change in overall Fos expression in rats recalling fear 48 hours after conditioning. Several factors may explain the discrepancy, including the fact prior work tested fear to the discrete cue in the same context in which conditioning occurred, whereas in our experiments testing took place in a novel chamber. Additionally, we sacrificed our animals 60 minutes following testing, while prior work used a 90-min time point. Finally, in the prior studies testing took place in a chamber in which the animals were able to make appetitive responses, whereas in our work no appetitive responses were made while the rats were tested. While this allows for a measure of conditioned suppression, there is evidence that allowing animals the ability to bar press for food while simultaneously testing them for fear to the cue produces a motivational conflict (i.e. fear vs reward) and that this is the key factor driving the engagement of PVT [45,46].

The basolateral amygdala is known to be involved in the acquisition of fear extinction [47,48], and there is evidence that BLA projections to the IL are also involved in this process [21]. However, whether the BLA and its connections participate in the recall of extinction is less clear. Imaging studies [21,49] have reported increased activity in BLA in animals recalling extinction memory. However, our prior work showed no difference in activation of the BLA in good versus poor extinction rats [7], and our results here show that extinction recall does not drive activation in the BLA broadly or within BLA projections to the IL. Consistent with our findings, whereas circuit manipulation studies indicate that IL inputs to the BLA are important for extinction learning [50,19], they are not necessary for extinction recall (Bukalo et al., 2016). However, a role for the BLA cannot be completely dismissed as recent evidence indicates that certain cell types in the BLA are required for extinction recall [51].

Finally, we found evidence that successful extinction recall involves the ventral hippocampus. This was specific to the ‘middle’ vHPC, as the same pattern was not observed in posterior regions. Consistent with prior work [52], we did not detect changes in Fos activation in vHPC afferents to the IL. There is considerable evidence that the renewal of fear seen when the CS is encountered outside the context in which extinction occurs requires the vHPC [53,54,49] and that it depends at least partially on vHPC inputs to the IL [11]. Based on these prior findings, we might have expected that poor extinction would be associated with increased activity in vHPC projections to the IL. However, this was not the case as there was no difference in Fos activity either in retrograde-labeled vHPC projections to IL or in non-labeled cells in the vHPC. This suggests that the failure to recall extinction in the extinction context may recruit different mechanisms than the renewal of fear.

The findings reported here help us to better understand how individual differences in extinction recall are reflected in differences in neural circuit activity. Our results might be relevant to post-traumatic stress disorder, which is known to involve excessive fear and a failure to extinguish fear responses. We showed that differences in extinction recall were associated with differences in neural activity both within, and outside, of projections to the IL. These differences were organized in discrete regions along the anteroposterior axis, further highlighting the importance of assessing brain function at the sub-region level. Shortcomings of the current approach include the correlational nature of the study and a focus on male rodents. Future studies should determine the neurobiological mechanisms of extinction learning in female rodents and employ approaches which allow for causal inferences.

## Acknowledgements

Kehinde Cole helped perform the experiments.

## References

1. Pavlov, I.P. Conditional reflexes: an investigation of the physiological activity of the cerebral cortex. (Oxford University Press, 1927).

2. Rothbaum, B.O., & Davis, M. Applying learning principles to the treatment of posttrauma reactions. Ann. N.Y. Acad. Sci. 1008(1), 112–121 (2003).

3. Milad, M.R. et al. Presence and acquired origin of reduced recall for fear extinction in PTSD: Results of a twin study. J. Psychiatr. Res. 42(7), 515–520 (2008).

4. Milad, M.R. et al. Neurobiological basis of failure to recall extinction memory in posttraumatic stress disorder. Biol. Psychiatry. 66(12), 1075–1082 (2009).

5. Bush, D.E.A., Sotres-Bayon, F., & LeDoux, J.E. Individual differences in fear: isolating fear reactivity and fear recovery phenotypes. J. Trauma. Stress. 20(4), 413–422 (2007).

6. Russo, A.S., & Parsons, R.G. Acoustic startle response in rats predicts inter-individual variation in fear extinction. Neurobiol. of Learn. and Mem. 139, 157–164 (2017).

7. Russo, A.S., Lee, J., & Parsons, R.G. Individual variability in the recall of fear extinction is associated with phosphorylation of mitogen-activated protein kinase in the infralimbic cortex. Psychopharmacology. 236(7), 2039–2048 (2019).

8. Yehuda, R., & LeDoux, J.E. Response variation following trauma: A translational neuroscience approach to understanding PTSD. Neuron. 56(1), 19–32 (2007).

9. Laurent, V., & Westbrook, R.F. Inactivation of the infralimbic but not the prelimbic cortex impairs consolidation and retrieval of fear extinction. Learn. Mem. 9, 520–529 (2009).

10. Kim, H., Cho, H., Augustine, G. J., & Han, J. Selective control of fear expression by optogenetic manipulation of infralimbic cortex after extinction. Neuropsychopharmacology. 41(5), 1261–1273 (2016).

11. Marek, R. et al. Hippocampus-driven feed-forward inhibition of the prefrontal cortex mediates relapse of extinguished fear. Nat. Neurosci. 21(3), 384–392 (2018).

12. Do-Monte, F.H., Manzano-Nieves, G., Quinõnes-Laracuente, K., Ramos-Medina, L., & Quirk, G.J. Revisiting the role of infralimbic cortex in fear extinction with optogenetics. J. Neurosci. 35(8), 3607–3615 (2015).

13. Hefner, K. et al. Impaired fear extinction learning and cortico-amygdala circuit abnormalities in a common genetic mouse strain. J. Neurosci. 28(32), 8074–8085 (2008).

14. Herry, C. & Mons, N. Resistance to extinction is associated with impaired immediate early gene induction in medial prefrontal cortex and amygdala. Eur. J. Neurosci. 20(2), 781–790 (2004).

15. Milad, M. R., & Quirk, G. J. Neurons in medial prefrontal cortex signal memory for fear extinction. Nature. 420(6911), 70–74 (2002).

16. Hoover, WB., & Vertes, R.P. Anatomical analysis of afferent projections to the medial prefrontal cortex in the rat. Brain Struct Funct. 212(2), 149–179 (2007).

17. Vertes, R.P. Differential projections of the infralimbic and prelimbic cortex in the rat. Synapse. 51, 32–58 (2003).

18. Adhikari, A. et al. Basomedial amygdala mediates top-down control of anxiety and fear. Nature. 527(7577), 179–185 (2015).

19. Bloodgood, D.W., Sugam, J.A., Holmes, A., & Kash, T.L. Fear extinction requires infralimbic cortex projections to the basolateral amygdala. Transl. Psychiatry. 8(1), 1–11 (2018).

20. Cho, J., Deisseroth, K., & Bolshakov, V.Y. Synaptic encoding of fear extinction in mPFC-amygdala circuits. Neuron. 80(6), 1491–1507 (2013).

21. Senn, V. et al. Long-range connectivity defines behavioral specificity of amygdala neurons. Neuron. 81(2), 428–437 (2014).

22. Jin, J. & Maren, S. Fear renewal preferentially activates ventral hippocampal neurons projecting to both amygdala and prefrontal cortex in rats. Sci. Rep. 5(1), 1–6 (2015).

23. Qin, C. et al. Dorsal hippocampus to infralimbic cortex circuit is essential for the recall of extinction memory. Cereb. Cortex. 31(3), 1707–1718 (2021).

24. Ramanathan, K.R., Jin, J., Giustino, T.F., Payne, M.R., & Maren, S. Prefrontal projections to the thalamic nucleus reuniens mediate fear extinction. Nat. Commun. 9, 4527; https://doi.org/10.1038/s41467-018-06970-z (2018).

25. Tervo, T.G.R. et al. A designer AAV variant permits efficient retrograde access to projection neurons. Neuron. 92(2), 372–382 (2016).

26. Tao, Y. et al. Projections from infralimbic cortex to paraventricular thalamus mediate fear extinction retrieval. Neurosci. Bull. 37, 229–241 (2020).

27. Li, S., & Kirouac, G. J. Sources of inputs to the anterior and posterior aspects of the paraventricular nucleus of the thalamus. Brain. Struct. and Funct. 217(2), 257–273 (2012).

28. Boorman, D.C., Brown, R., & Keay, K.A. Periaqueductal gray inputs to the paraventricular nucleus of the thalamus: Columnar topography of glucocorticoid (in)sensitivity. Brain. Res. 1750; https://doi.org/10.1016/j.brainres.2020.147171 (2021).

29. McNally, G.P., Pigg, M., & Weidemann, G. Opioid receptors in the midbrain periaqueductal gray regulate extinction of Pavlovian fear conditioning. J. Neurosci. 24(31), 6912–6919 (2004).

30. McNally, G. P., Lee, B.-W., Chiem, J. Y., & Choi, E. A. The midbrain periaqueductal gray and fear extinction: Opioid receptor subtype and roles of cyclic AMP, protein kinase A, and mitogen-activated protein kinase. Behav. Neurosci. 119(4), 1023–1033 (2005).

31. Parsons, R.G., Gafford, G.M., & Helmstetter, F.J. Regulation of extinction-related plasticity by opioid receptors in the ventrolateral periaqueductal gray matter. Frontiers in Behavioral Neuroscience. 4(44); https://doi.org/10.3389/fnbeh.2010.00044 (2010).

32. Watson, T.C., Cerminara, N.L., Lumb, B.M., & Apps, R. Neural correlates of fear in the periaqueductal gray. J. Neurosci. 36(50), 12707–12719 (2016).

33. McGinty, J.F. & Otis, J.M. Heterogeneity in the paraventricular thalamus: The traffic light of motivated behaviors. Front. Behav. Neurosci. 181; https://doi.org/10.3389/fnbeh.2020.590528 (2020).

34. Gao, C. et al. Two genetically, anatomically and functionally distinct cell types segregate across the anteroposterior axis of paraventricular thalamus. Nat. Neurosci. 23(2), 217–228 (2020).

35. Li, S. & Kirouac, G.J. Projections from the paraventricular nucleus of the thalamus to the forebrain, with special emphasis on the extended amygdala. J. Comp. Neurol. 506(2), 263–287 (2008).

36. Mathur, B.N. The claustrum in review. Front. Syst. Neurosci. 8, 1–11 (2014).

37. Goll, Y., Atlan, G., & Citri, A. Attention : The claustrum. Trends Neurosci. 38(8), 486–495 (2015).

38. Luppi, P.H., Billwiller, F., & Fort, P. Selective activation of a few limbic structures during paradoxical (REM) sleep by the claustrum and the supramammillary nucleus: evidence and function. Curr. Opin. Neurobiol. 44, 59–64 (2017).

39. Jackson, J., Karnani, M.M., Zemelman, B.V, Burdakov, D., & Lee, A K. Inhibitory control of prefrontal cortex by the claustrum. Neuron. 99(5), 1029–1039 (2018).

40. Beck, C. H., & Fibiger, H. C. Conditioned fear-induced changes in behavior and in the expression of the immediate early gene c-fos: with and without diazepam pretreatment. J. Neurosci. 15(1), 709–720 (1995).

41. Kitanishi, T., & Matsuo, N. Organization of the claustrum-to-entorhinal cortical connection in mice. J. Neurosci. 37(2), 269–280 (2017).

42. Do-Monte, F.H., Quinõnes-Laracuente, K., & Quirk, G.J. A temporal shift in the circuits mediating retrieval of fear memory. Nature. 519, 460–463 (2015).

43. Li, Y., Dong, X., Li, S., & Kirouac, G. J. Lesions of the posterior paraventricular nucleus of the thalamus attenuate fear expression. Front. Behav. Neurosci. 8(94); https://doi.org/10.3389/fnbeh.2014.00094 (2014).

44. Padilla-Coreano, N., Do-Monte, F.H., & Quirk, G.J. A time-dependent role of midline thalamic nuclei in the retrieval of fear memory. Neuropharmacology. 62(1), 457–463 (2012).

45. Choi, E. A., & McNally, G. P. Paraventricular thalamus balances danger and reward. J. Neurosci. 37(11), 3018–3029 (2017).

46. Choi, E. A., Jean-Richard-dit-Bressel, P., Clifford, C. W., & McNally, G. P. Paraventricular thalamus controls behavior during motivational conflict. J. Neurosci. 39(25), 4945–4958 (2019).

47. Falls, W. A., Miserendino, M. J., & Davis, M. Extinction of fear-potentiated startle: blockade by infusion of an NMDA antagonist into the amygdala. J. Neurosci. 12(3), 854–863 (1992).

48. Sierra-Mercado, D., Padilla-Coreano, N., & Quirk, G.J. Dissociable roles of prelimbic and infralimbic cortices, ventral hippocampus, and basolateral amygdala in the expression and extinction of conditioned fear. Neuropsychopharmacology. 36(2), 529–538 (2011).

49. Knapska, E., & Maren, S. Reciprocal patterns of c-Fos expression in the medial prefrontal cortex and amygdala after extinction and renewal of conditioned fear. Learn. Mem. 734(16), 486–493 (2009).

50. Bukalo, O. et al. Prefrontal inputs to the amygdala instruct extinction memory formation. Sci. Adv. 1(6); doi: 10.1126/sciadv.1500251

51. Zhang, X., Kim, J., & Tonegawa, S. Amygdala reward neurons form and store fear extinction memory. Neuron. 105(6), 1077–1093 (2020).

52. Wang, Q., Jin, J., & Maren, S. Renewal of extinguished fear activates ventral hippocampal neurons projecting to the prelimbic and infralimbic cortices in rats. Neurobiol. Learn. Mem. 134, 38–43 (2016).

53. Hobin, J. A., Ji, J., & Maren, S. Ventral hippocampal muscimol disrupts context-specific fear memory retrieval after extinction in rats. Hippocampus. 16(2), 174–182 (2006).

54. Orsini, C. A., Kim, J. H., Knapska, E., & Maren, S. Hippocampal and prefrontal projections to the basal amygdala mediate contextual regulation of fear after extinction. J. Neurosci. 31(47), 17269–17277 (2011).

